# Two Novel Red-FRET ERK Biosensors in the 670-720nm Range

**DOI:** 10.1101/2024.11.30.626109

**Authors:** Nicholaus L. DeCuzzi, Jason Y. Hu, Florene Xu, Brayant Rodriguez, Michael Pargett, John G. Albeck

## Abstract

Cell fate decisions are regulated by intricate signaling networks, with Extracellular signal-Regulated Kinase (ERK) being a central regulator. However, ERK is rarely the only signaling pathway involved, creating a need to study multiple signaling pathways simultaneously at the single-cell level. Many existing fluorescent biosensors for ERK and other pathways have significant spectral overlap, limiting their ability to be multiplexed. To address this limitation, we developed two novel red-FRET ERK biosensors, REKAR67 and REKAR76, which operate in the 670-720 nm range using miRFP670nano3 and miRFP720. REKAR67 and REKAR76 differ in fluorophore position, which impacts biosensor characteristics; REKAR67 displayed a higher dynamic range but greater signal variance than REKAR76. Mixed populations of REKAR67 or REKAR76 displayed similar Signal-to-Noise ratio (SNR), but in clonal cell populations, REKAR76 had a significantly higher SNR. Overall, our red-FRET ERK biosensors were highly consistent with existing ERK FRET biosensors and in reporting ERK activity and are spectrally compatible with CFP/YFP FRET and cpGFP -based biosensors. Both REKAR biosensors expand the available methods for measuring single-cell ERK activity.

## Introduction

Extracellular-signal Regulated Kinase (ERK) is a terminal kinase of the Mitogen Activated Protein Kinase (MAPK) signaling pathway and is a key regulator of cell fates including proliferation, growth, and apoptosis [1,2]. These cell fate decisions are determined by the temporal activity patterns of ERK, motivating the need to quantify ERK activity at the single-cell level. Fluorescent biosensors have revealed significant insight into the signaling activity of ERK, providing real-time activity readouts with little perturbation to normal kinase function [3]. For example, ERK activity often occurs in radiating waves originating at discrete points, both *in vivo* [4] and in various cell culture models [5–8]. These waves have functional significance in maintaining homeostasis of epithelial layers. Furthermore, oncogenic mutations in RAS or EGFR modulate the forms of ERK kinetics [9,10], impacting the expression of downstream genes [11]. A key area of interest is to now understand how dynamic ERK activity patterns integrate with other signaling pathways. However, the currently available biosensors have limitations in specificity and fluorescence properties that create significant difficulties for multi-biosensor experiments.

One widely used design for ERK biosensors uses Förster Resonance Energy Transfer (FRET) between two coupled fluorophores, where the excitation energy of one (FRET donor) is transferred to and subsequently emitted by the second fluorophore (FRET acceptor), causing a ratiometric increase in acceptor emission. ERK Kinase Activity Reporters (EKARs) are a class of biosensors that use FRET, typically between cyan and yellow fluorescent proteins (CFP and YFP), to quantify ERK activity in living cells. For example, the EKAR-EV biosensor construct contains the CFP variant ECFP tethered to the YFP variant YPet via a flexible linker. Within this linker region, there is a ERK substrate recognition domain containing a phospho-acceptor threonine residue, and a phospho-recognition domain. When the threonine residue within the ERK substrate domain is phosphorylated, it is then recognized and bound by the phospho-amino acid-binding WW domain. The intramolecular interaction of the phosphorylated residue with the WW domain brings the two fluorophores into close proximity, increasing the measured FRET.

There have been multiple improvements to EKAR since its initial inception [12], including changes to the linker length [13], usage of brighter and more pH-stable CFP and YFP variants [14], and changes in FP position. Recent iterations of EKAR, EKAREN4 and EKAREN5, addressed the non-specific phosphorylation of the biosensor by CDK1, which causes non-ERK specific signals in G2/M phases of the cell cycle [15]. Two amino acid substitutions in the substrate domain were sufficient to nearly eliminate the CDK-mediated response of the biosensor. By making changes in linker length, EKAREN4 and EKAREN5 were further modified to have different dynamic ranges relative to ERK activity, with EKAREN4 showing a larger range overall and higher saturation point. EKAREN5 has a greater degree of sensitivity and is best for detecting low levels of ERK activity [15].

While CFP/YFP FRET-based ERK biosensors offer valuable insights into ERK activity, their spectral overlap with many other biosensors limits the ability to simultaneously monitor multiple signaling pathways, which is crucial for understanding complex cell fate decisions. For example, measuring ERK and Protein Kinase B (AKT) activity together has enabled insights into their combined roles in promoting cell proliferation or differentiation [7,16]. Such studies have relied on translocation-based reporters for ERK and AKT, which do not provide spatial resolution, and could be enhanced using reporters such as EKAREN4/5 and AKT cpGFP-based biosensor, ExRai-AKTAR2 [17]. However, these biosensors have overlapping spectral properties, making multiplexing at the single-cell level challenging. The development of spectrally distinct fluorescent biosensors would alleviate this issue and allow for multiple signaling pathways to be measured in the same cell. Efforts to develop red/far-red FRET versions of CFP/YFP FRET biosensors have been limited, and in particular, the influence of FP location has not been investigated despite being previously demonstrated to affect the dynamic range of CFP/YFP FRET biosensors.

Here, we addressed the wavelength limitation in EKAR biosensors by developing two red-shifted versions of EKAREN4, designated as REKAR67 and REKAR76. These new constructs operate in the 670-720 nm range, using the fluorophores miRFP670nano3 and miRFP720 as FRET donor and acceptor, respectively. The two versions of REKAR were created with alternate ordering of the two red FPs to determine if FP position impacts biosensor characteristics, including dynamic range, noise variance, and Signal to Noise Ratio (SNR). We found that of the two, REKAR67 shows a better dynamic range while REKAR76 has reduced variance. We compared REKAR variants directly to EKAREN4 in dual biosensor-expressing cells, finding that both REKAR versions are comparable to the original EKAREN4 in responses to Epidermal Growth Factor (EGF) stimulation.

## Materials and Methods

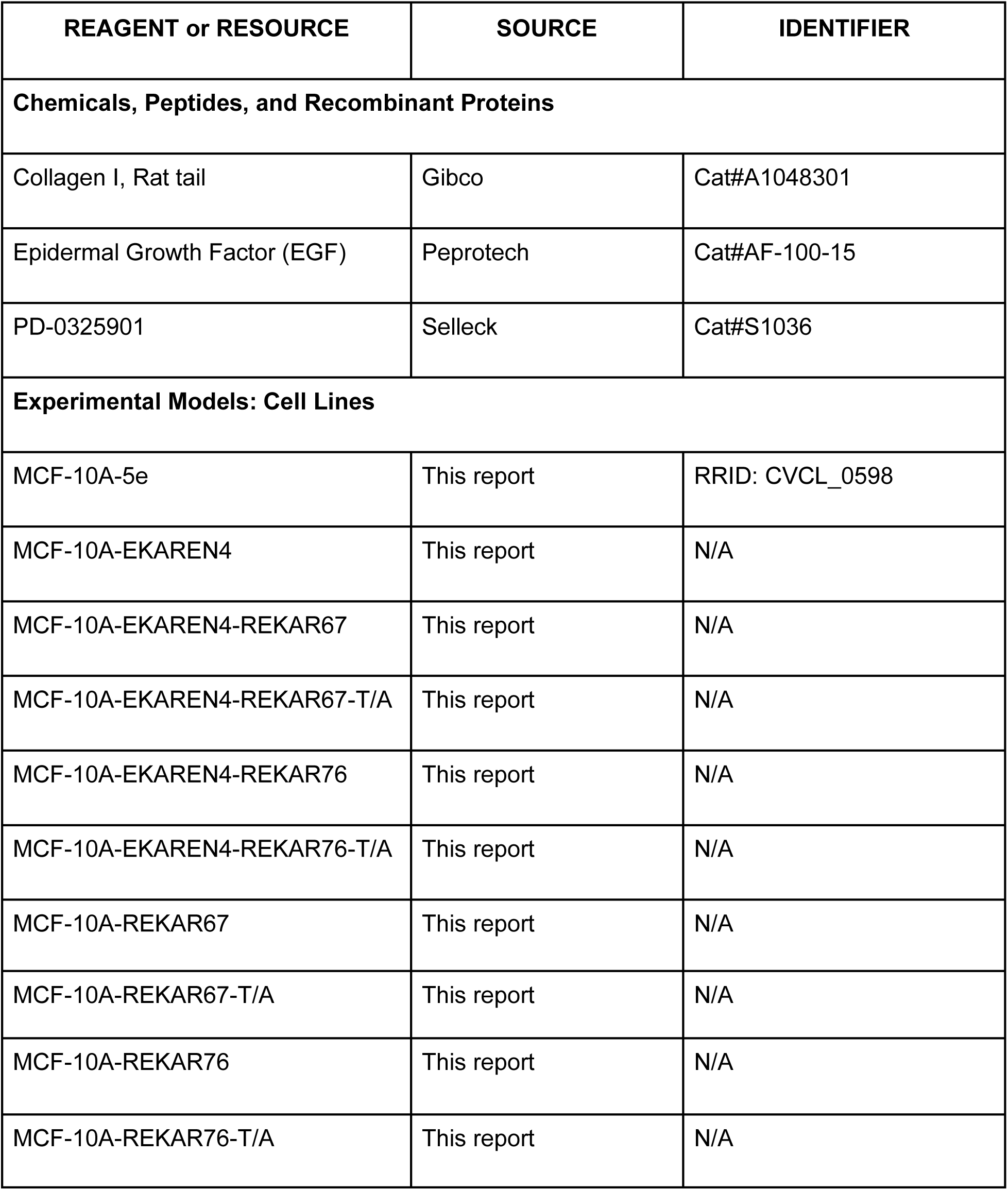
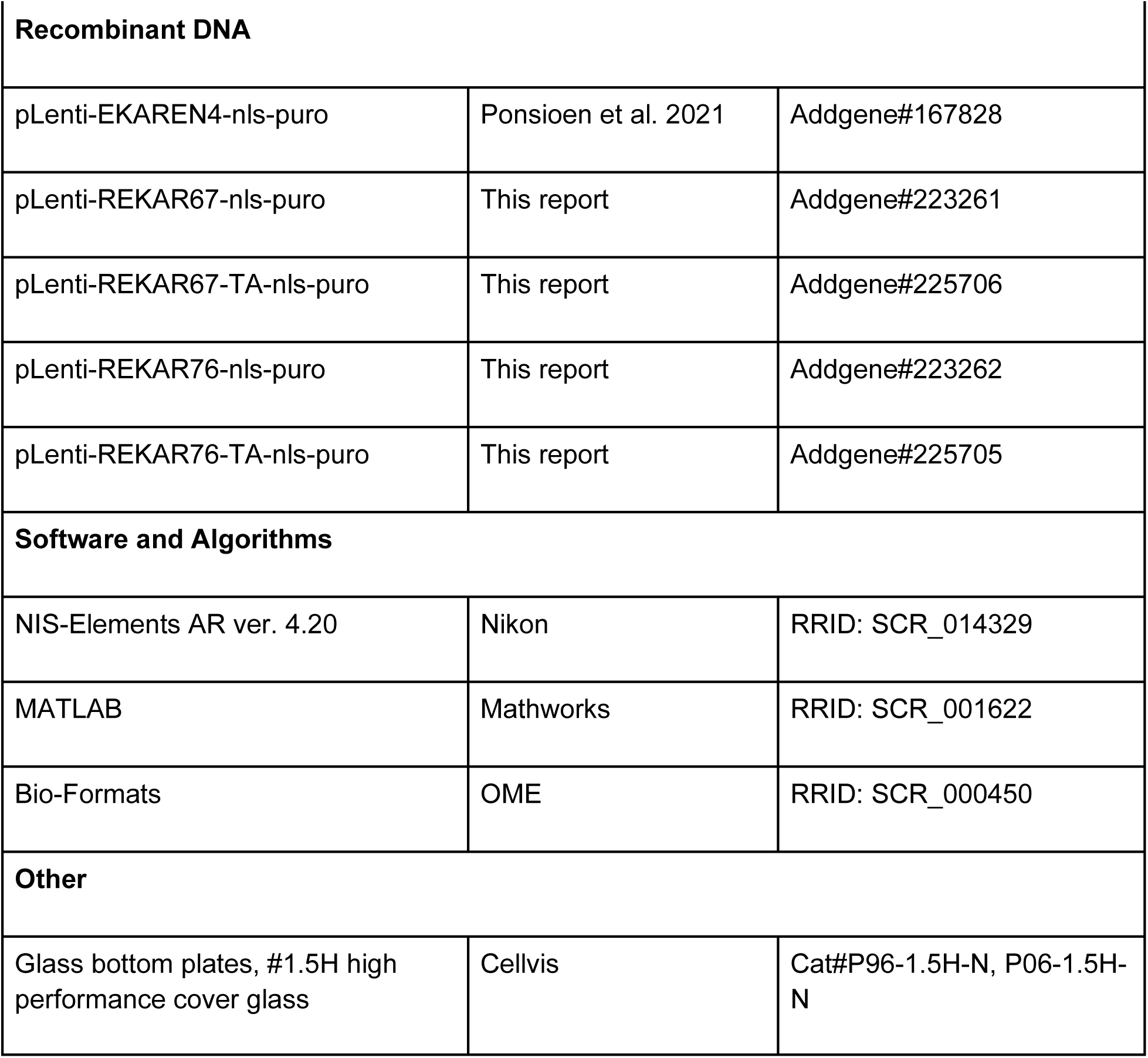
Key Resources Table

| REAGENT or RESOURCE | SOURCE | IDENTIFIER |
| --- | --- | --- |
| **Chemicals, Peptides, and Recombinant Proteins** |  |  |
| Collagen I, Rat tail | Gibco | Cat#A1048301 |
| Epidermal Growth Factor (EGF) | Peprotech | Cat#AF-100-15 |
| PD-0325901 | Selleck | Cat#S1036 |
| **Experimental Models: Cell Lines** |  |  |
| MCF-10A-5e | This report | RRID: CVCL_0598 |
| MCF-10A-EKAREN4 | This report | N/A |
| MCF-10A-EKAREN4-REKAR67 | This report | N/A |
| MCF-10A-EKAREN4-REKAR67-T/A | This report | N/A |
| MCF-10A-EKAREN4-REKAR76 | This report | N/A |
| MCF-10A-EKAREN4-REKAR76-T/A | This report | N/A |
| MCF-10A-REKAR67 | This report | N/A |
| MCF-10A-REKAR67-T/A | This report | N/A |
| MCF-10A-REKAR76 | This report | N/A |
| MCF-10A-REKAR76-T/A | This report | N/A |
| **Recombinant DNA** |  |  |
| pLenti-EKAREN4-nls-puro | Ponsioen et al. 2021 | Addgene#167828 |
| pLenti-REKAR67-nls-puro | This report | Addgene#223261 |
| pLenti-REKAR67-TA-nls-puro | This report | Addgene#225706 |
| pLenti-REKAR76-nls-puro | This report | Addgene#223262 |
| pLenti-REKAR76-TA-nls-puro | This report | Addgene#225705 |
| **Software and Algorithms** |  |  |
| NIS-Elements AR ver. 4.20 | Nikon | RRID: SCR_014329 |
| MATLAB | Mathworks | RRID: SCR_001622 |
| Bio-Formats | OME | RRID: SCR_000450 |
| **Other** |  |  |
| Glass bottom plates, #1.5H high performance cover glass | Cellvis | Cat#P96-1.5H-N, P06-1.5H-N |

## Resource Availability

Further information and requests for resources and reagents should be directed to and will be fulfilled by the Lead Contact Dr. John Albeck.

## Materials and Availability

REKAR67, REKAR67-T/A, REKAR76, and REKAR76-T/A plasmids are available from Addgene (plasmid numbers: 223261, 225706, 223262, and 225705, respectively). Cell lines can be provided upon request to Dr. John Albeck.

## Data and Code Availability

Image data files collected via NIS elements are available upon request, but could not be uploaded due to large file size (50-500 GB per ND2 file).

All data processing was performed in MATLAB using previously described methods.[10,18,19] Code and methods available on a github repository: https://github.com/Albeck-Lab/REKAR

Experimental Model and Subject Details:

## Cell Culture and Media

Human mammary epithelial cells, MCF-10A clone 5E, were cultured in MCF-10A Growth Medium (see Media Composition Table) as described in [20]. Low passage stocks from the parental MCF-10A-5E clone were used to create REKAR67, REKAR67-T/A, REKAR76, and REKAR76-T/A containing cell lines. An MCF-10A clone that expresses EKAREN4 was used to create EKAREN4-REKAR67, EKAREN4-REKAR67-T/A, EKAREN4-REKAR76, and EKAREN4-REKAR76-T/A cell lines. All cell lines were passaged before reaching 85% confluency.

## Biosensor construction

REKAR67 and REKAR76 plasmids were assembled by replacing the YFP and CFP fluorophores of pLJM1-EKAREN4 with miRFP670nano3 and miRFP720 from synthesized fragments via Gibson assembly (NEB M5510A). REKAR67 is defined by the N-terminal location of miRFP670nano3, miRFP720. REKAR76 is characterized by the N-terminal location of miRFP720 followed by miRFP670nano3. The negative controls, REKAR67-T/A and REKAR76-T/A, were generated with a T498A mutation, preventing phospho-recognition of the substrate domain.

## Biosensor delivery

The biosensors REKAR67 or REKAR76 were transduced into MCF-10A-5E cells, or into MCF-10A-5E cells with a clonal integration of EKAREN4, via lentiviral transduction with pLenti-REKAR viral particles [21]. Lentiviral particles were synthesized by transfecting HEK 293T cells with the Fugene HD transfection reagent (SKU: HD-1000). Lentivirus transduction was carried out in 8 μg/mL polybrene treated media with centrifugation at 100 x g for 30 minutes. Flow cytometry sorting was used to isolate biosensor positive cells with a Beckman Coulter MoFlo Astrios EQ cell sorter (UC Davis Flow Cytometry Shared Resource).

## Live-cell microscopy experiments

Live-cell imaging experiments were conducted following previously described protocols [18,19]. MCF-10A cells were plated on 96-well imaging plates (Cellvis #:P96-1.5H-N) at least 48 hours prior to experiments. Cells were seeded in MCF-10A Growth Medium supplemented with 25 μM biliverdin, then transferred to MCF-10A Imaging Medium at least 16 hours prior to imaging (see Media Composition tables). During live-cell imaging experiments, cells were maintained at 37°C with 5% CO_2_. Images of cells were taken every 6 minutes using a Teledyne Photometrix Kinetix sCMOS camera and a Nikon (Tokyo, Japan) 20X/0.75 NA Plan Apo objective on a Nikon Eclipse Ti2 inverted microscope, equipped with a Lumencor SPECTRA III light engine. Experiments were conducted using the following fluorescence filter sets: CFP (#49001, Chroma), YFP (#49003, Chroma), miRFP670 (custom T647lpxr – ET667/30m, Chroma, with 637nm laser excitation) and FRET720 (custom T647lpxr – FF01-730/39-32, Chroma and Semrock, with 637nm laser excitation). EKAREN4 measurements were made using CFP and YFP filters, and REKAR measurements were made using miRFP670 and FRET720 filters.

## Biosensor Quantification

The EKAREN4 biosensor was quantified as described in [18]. Briefly, the response of the EKAREN4 biosensor is represented by the value *Ef_A_*, which represents the product of *E*, the FRET efficiency of the fluorophore pair in the associated configuration, and *f_A_*, the fraction of fluorophore pairs in that associated configuration. This value is computed as: *Ef_A_* = 1 − 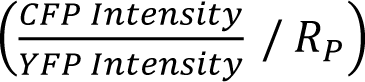, where *R_P_* is the ratio of total channel gain in the CFP channel over that in the YFP channel. Each gain is computed as the spectral product of the contributing imaging system parameters: the relative excitation intensity, exposure time, molar extinction coefficient, quantum yield, light source spectrum, filter transmissivities, and fluorophore absorption and emission spectra.

The response of the REKAR biosensor was determined in the same fashion. However, given the spectral overlap of the miRFP670nano3 and miRFP720 fluorophores, it is not feasible to measure their contributions without significant spectral overlap. Accordingly, a more general form of the FRET measurement model was necessary to account for spectral cross-talk terms. Working from the imaging system model presented in the “FRET reporter measurement and correction” section of the appendix to [14] (eqns. 1-3), we model the intensity ratio of the FRET720 imaging channel to the RFP670 channel as:

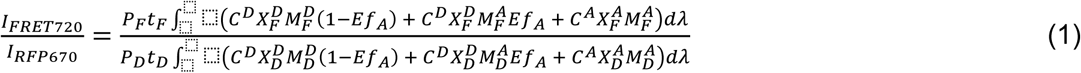

Herein, *P* denotes the relative power delivered, *t* the exposure time, *C* the concentration of a fluorophore, *X* an excitation spectrum (the spectral product of the light source spectrum, the excitation filter, and the fluorophore absorption spectrum), and *M* an emission spectrum (product of the fluorophore quantum yield, emission spectrum and the emission filter transmissivity). In superscripts, D refers to the Donor fluorophore (miRFP670nano3), and A the acceptor (miRFP720). In subscripts, D refers to the Donor imaging channel (that using the RFP670 filter set), and F the FRET imaging channel (via the FRET720 filter set).

In this case, the spectral cross-talk term for the Donor emission, *^A^_D_* (the fraction of light emitted by the Acceptor that is collected, relative to that collected from the Donor), is estimated to be on the order of 0.1 or less, confirmed by the absence of signal in the RFP670 channel from isolated miRFP720 molecules (data not shown). This allows simplification of the denominator of the spectra ratio model by neglecting this term:

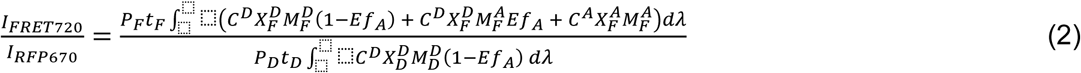

This model may then be rearranged, employing several substitutions for convenience. Define the normalized intensity ratio (NIR), and spectral ratios for cross-talk (XT), channel, and FRET:

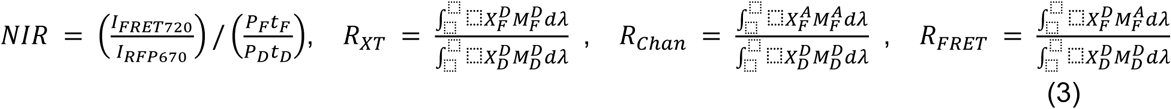

The final model may then be written as:

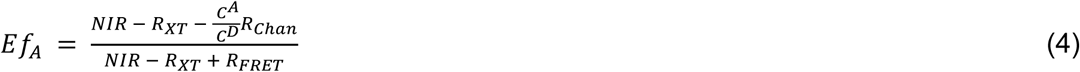

Each of the three spectral “ratios” may be computed via calibrations of the imaging system. Because the crosstalk ratio (*R_XT_*) is largely composed of effects on the edges/tails of spectra, it may be necessary to make a direct calibration measurement of the interaction. This necessity can arise from minor variance in the spectra provided by light source and filter manufacturers, as well as those reported for the fluorophores. At the edges/tails, where spectral power or transmissivity is changing rapidly with wavelength, minor errors in different spectra may stack to yield significant errors in the computed spectral products and ratios. By imaging the miRFP670nano3 fluorophore alone via both the RFP670 and FRET720 filter sets, the crosstalk ratio was directly measured, with a value of 0.578 on our hardware (data not shown), which is less than 5% greater than the computed value of 0.551.

## Single-cell data processing

Single-cell values for the fractions of active EKAREN4 and/or REKAR biosensors were quantified, tracked over time, and filtered to omit data tracks that did not last >85% of the movie length or that showed signals outside of expected ranges (EKAREN4 = 0.3 - 0.65 & all REKAR variants = 0.15 - 0.4). Data were normalized (where indicated) at the single-cell level by subtracting the cell’s minimum signal and dividing that signal by its own maximum signal (after min subtraction), to give a signal range for each reporter of 0 to 1 to every cell. Data normalization was performed on the EKAREN4, REKAR67, and REKAR76 biosensor signals of all cells as indicated.

### Media Composition Tables

**Table 1.** MCF-10A Growth Medium (GM)

| Reagent | Supplier | Item # | Stock Concentrations | Final Concentrations |
| --- | --- | --- | --- | --- |
| DMEM/F12 | Gibco | 11320033 | 1X | 1X |
| Horse Serum | Gibco | 26050088 | 100 % v/v | 5% v/v |
| Cholera Toxin | Sigma | C8052 | 1 mg/mL | 100 ng/mL |
| EGF | Peprotech | AF-100-15 | 100 µg/mL | 5 ng/mL |
| Hydrocortisone | Sigma | H0888 | 1 mg/mL | 0.5 µg/mL |
| Insulin | Sigma | I9278 | 10 mg/mL | 10 µg/mL |
| Biliverdin* | Sigma | 30891 | 32 mM | 25 µM |

\*Only supplemented into growth medium 24-48 hours prior to use for an imaging experiment.

**Table 2.** MCF-10A Imaging Medium (IM)

| Reagent | Supplier | Item # | Stock Concentrations | Final Concentrations |
| --- | --- | --- | --- | --- |
| Imaging base-DMEM/F12* | UC Davis Veterinary Medicine Biological Media Services |  | 1X | 1X |
| Bovine Serum Albumin | Sigma | A7906 | 1X | 0.1% w/v |
| Glucose | Fisher |  | 1.74 M | 17.4 mM |
| Pyruvate | Gibco | 11360070 | 100 mM | 1 mM |
| Hydrocortisone | Sigma | H0888 | 1 mg/mL (2.76mM) | 0.5 ug/mL (1.38 μM) |
| Cholera Toxin | Sigma | C8052 | 1 mg/mL | 100 ng/mL |

*\* Custom F12/DMEM similar to GIBCO 11320033 lacking Glucose, Glutamine, Pyruvate, Folic Acid, Riboflavin, and Phenol Red.*

**Table 3.**
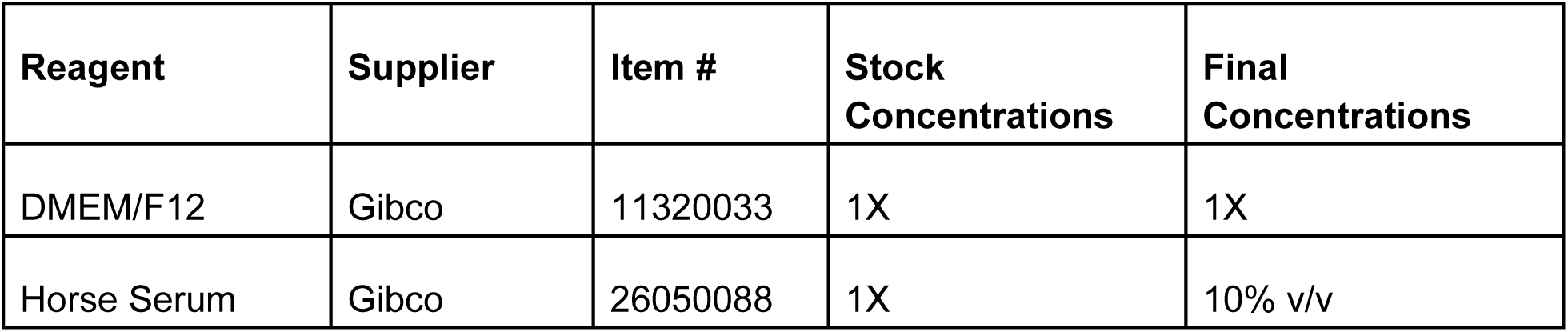
MCF-10A Resuspension Medium

| Reagent | Supplier | Item # | Stock Concentrations | Final Concentrations |
| --- | --- | --- | --- | --- |
| DMEM/F12 | Gibco | 11320033 | 1X | 1X |
| Horse Serum | Gibco | 26050088 | 1X | 10% v/v |

**Table 4.** Molecular Reagents

| Reagent | Supplier | Item # | Stock Concentrations | Final Concentrations |
| --- | --- | --- | --- | --- |
| Fugene® Hd Transfection Reagent | FuGene | HD-1000 | 1X | 6% w/v |
| Gibson Assembly Master Mix | New England BioLabs | M5510A | 2X | 1X |

## Results

### Red-far red-FRET ERK biosensors accurately measure changes in ERK activity

To develop a FRET-based ERK biosensor that does not overlap with fluorescence in the 400-600 nm range, we constructed REKAR67 and REKAR76 by replacing the Turquoise2 (CFP) and Ypet (YFP) fluorophores of EKAREN4 [15] with miRFP670nano3 [22] and miRFP720 [23] (**Figure 1A-C**). REKAR67 and REKAR76 were designed with alternate fluorophore positions to identify the best configuration for imaging properties such as dynamic range (minimum to maximum signal measured). Fluorophore position has been found to be important in optimizing ERK CFP/YFP FRET biosensors [24]. The two constructs were named based on the N or C terminal location of the red fluorescent proteins (N to C terminal order):

1. REKAR67: miRFP670nano3 - Phospho-recognition domain (WW) - linker domain - ERK substrate - miRFP720 (**Figure 1B**)
2. REKAR76: miRFP720 - Phospho-recognition domain (WW) - linker domain - ERK substrate - miRFP670nano3 (**Figure 1C**)

**Figure 1.**
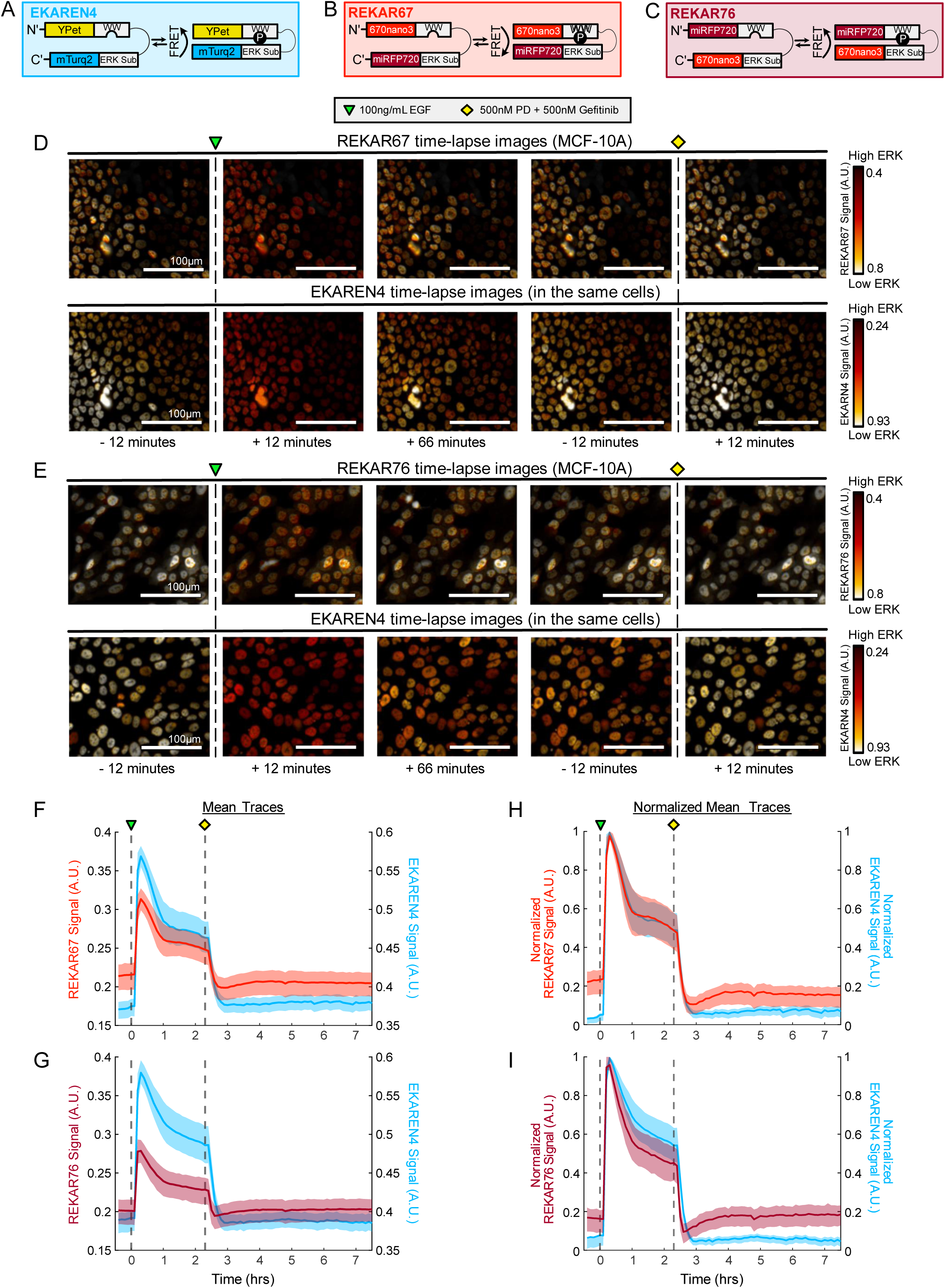
Red-FarRed ERK FRET-based biosensor designs and validation. **(A-C)** Graphical representations of the ERK biosensors EKAREN4, REKAR67, and REKAR76. **(D&E)** Pseudo-colored images of biosensor activity from cells co-expressing the REKAR and EKAREN4 biosensors. *Top row* shows REKAR67 or REKAR76 and *bottom row* shows EKAREN4. ERK activity corresponds to the indicated color scale (white = low ERK, red = high ERK). Images show before and after EGF stimulation then MEK/EGFR inhibition. Scale bars are 100μm. **(F-G)** Mean FRET signal of REKAR67 (red) or REKAR76 (maroon) overlaid on the mean EKAREN4 signal (blue) of the same cells. **(H-I)** Normalized data from **F&G**.

We validated REKAR67 and REKAR76 by stably expressing each of these biosensors in MCF-10A cells already expressing the EKAREN4 biosensor and comparing their responses to canonical ERK activators (100 ng/mL EGF) and pathway inhibition (500nM PD-0325901 and gefitinib) (**Figure 1D&E; Sup. Videos 1-4**). Treatment with EGF caused a sharp increase in EKAREN4, REKAR67, and REKAR76 signals, peaking approximately 30 minutes after treatment, whereas treatment with PD-0325901 and gefitinib resulted in a decrease to the minimum signal within 20 minutes (**Figure 1F&G**). To better compare REKAR67 and REKAR76 to EKAREN4 we normalized each ERK biosensor to its own minimum and maximum signal within the cell (**Figure 1H&I**). With this normalization, the rise and fall of both REKAR variants was highly similar to EKAR and to one another. However, a consistent difference between EKAR and both REKAR variants was that, following inhibitor treatment, REKAR levels first fell to a minimum lower than the initial baseline, and then recovered within about 60 minutes to a level similar to the baseline. In contrast, EKAR signals fell immediately to a constant level similar to the initial baseline. Additionally, we noticed that the 25th-75th interquartile range for both REKAR reporters tended to be broader than for EKAREN4 during the pre-stimulation and post-inhibition phases, indicating a greater degree of variation. Nonetheless, despite these differences, the similarities of REKAR67 and REKAR76 signals to EKAREN4 signals suggests both red-FRET biosensors also measure changes in ERK activity.

### REKAR67 and REKAR76 signal is dependent on ERK substrate phosphorylation status

We verified that neither configuration of red-FRET ERK biosensor acquired unintended off target phosphorylation sites by generating variants of both sensors in which the phosphoacceptor threonine of the ERK substrate sequence was substituted with alanine. This T/A mutation eliminates the ERK phosphorylation target within the biosensors, and these variants should not exhibit a change in FRET signal when any kinase, including ERK, is active within the cell. We generated cells expressing EKAREN4 and either REKAR67-T/A or REKAR76-T/A (**Sup. Figure 1A&B; Sup. Videos 5-8**). Unlike their functional counterparts, neither REKAR-T/A variant responded to EGF or PD-0325901 and gefitinib treatment, while the co-expressed EKAREN4 maintained its responsiveness to both treatments (**Sup. Figure 1 C&D**). Therefore, the response of both REKAR biosensors is strictly dependent on the phosphorylation of the threonine residue of the ERK substrate sequence. Furthermore, this experiment demonstrated definitively that changes in REKAR sensor signals are spectrally independent from changes occurring in a co-expressed CFP/YFP biosensor.

### REKAR67 and REKAR76 both accurately measure single-cell ERK activity

To further compare the REKAR variants to EKAREN4, we generated visualizations of individual cells, using a similar normalization scheme to the mean plots displayed in **Figure 1** (**Figure 2A&B**). In these visualizations, we again found a close concordance between REKAR signals and EKAREN4. Notably, when cells differed from one another in the shape of their EGF-induced ERK activity peak, this shape was shared by both the REKAR and EKAREN4 signals within the same cell. Thus, when used to read out cell-to-cell differences, EKAREN4 and REKAR provide similar information. However, we also noted that REKAR signals showed a greater degree of high-frequency variation, which was not obviously correlated with the EKAREN4 signal in the same cell nor from cell-to-cell. This observation suggests that our measurements of REKAR signals are more subject to noise than those of EKAREN4. To examine genetic variation (e.g., different reporter integration positions in the genome) as a potential source of noise, we derived clonal populations of cells carrying each REKAR variant. In their average responses, these clonal lines were highly similar to the clonal populations (**Figure 2 C&D**). While the interquartile range for the clones was smaller than the mixed populations, it was still larger than EKAR (**Figure 1 F-I**). Thus, there remains an additional source of noise in EKAR that we have not yet accounted for.

**Figure 2.**
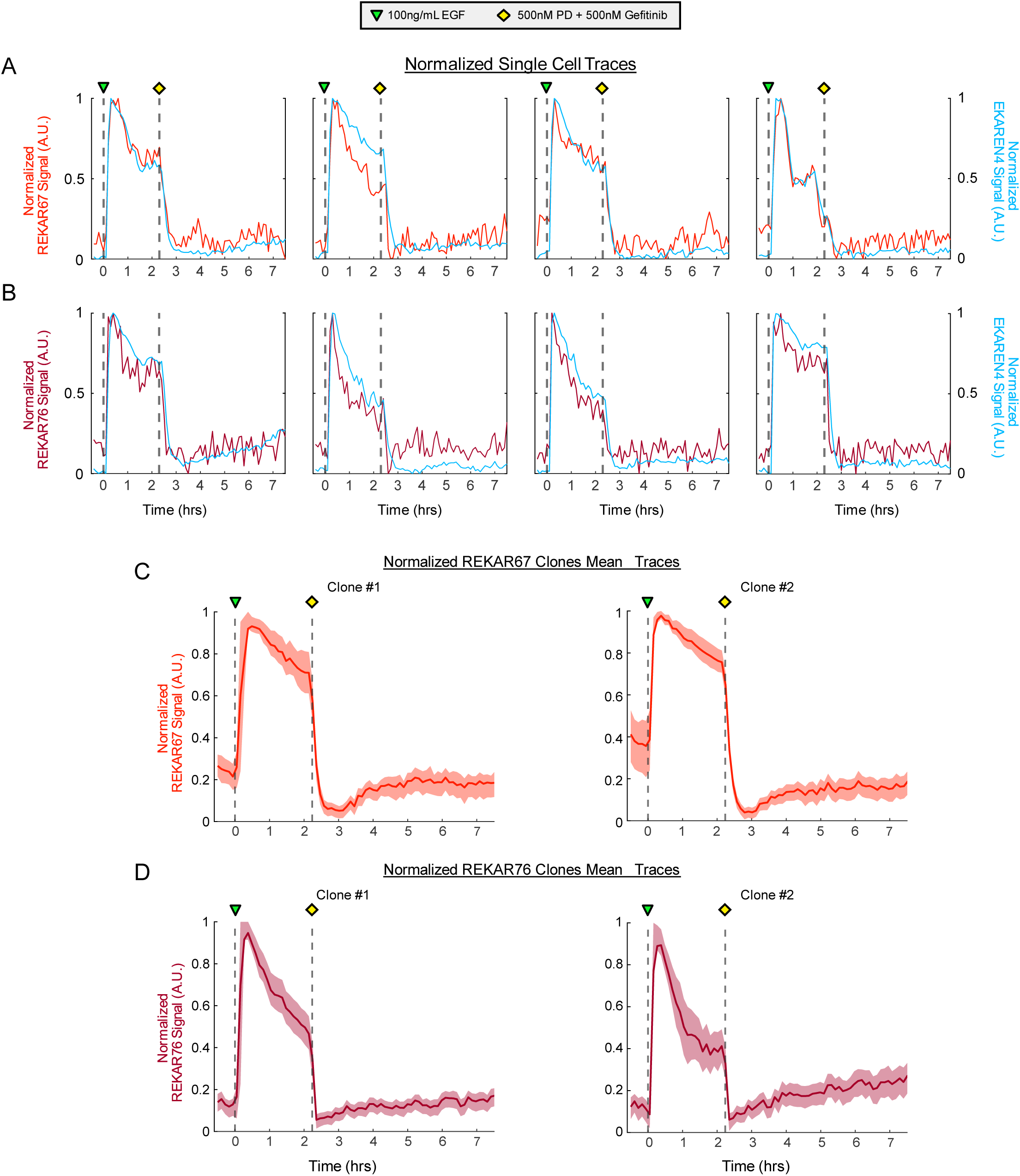
Single-cell REKAR-EKAREN4 and REKAR clone activity traces. **(A)** Normalized MCF10A REKAR67-EKAREN4 double expressing cell activity traces overlayed (red and blue, respectively). **(B)** Normalized MCF10A REKAR76-EKAREN4 double expressing cell activity traces overlayed (maroon and blue, respectively). **(C)** Normalized mean REKAR67 FRET activity of clonal REKAR67 expressing MCF10A cells. (**D)** Normalized mean REKAR76 FRET activity of clonal REKAR76 expressing MCF10A cells.

### Signal-to-noise properties of REKAR67 and REKAR76

We next evaluated and compared the signal-to-noise ratio (SNR) for REKAR67 and REKAR76. We defined “Signal” as the dynamic range of the biosensor, from maximal activation (following EGF stimulation) to maximal suppression of ERK (following a PD and gefitinib treatment). “Noise” was defined as the frame-to-frame variance of the signal following inhibition (**Figure 3A**), because we expected that true ERK activity would be fully suppressed under these conditions; the observation that the T/A variants exhibited similar Noise values confirms this expectation. We compared both mixed and clonal populations of cells.

**Figure 3.**
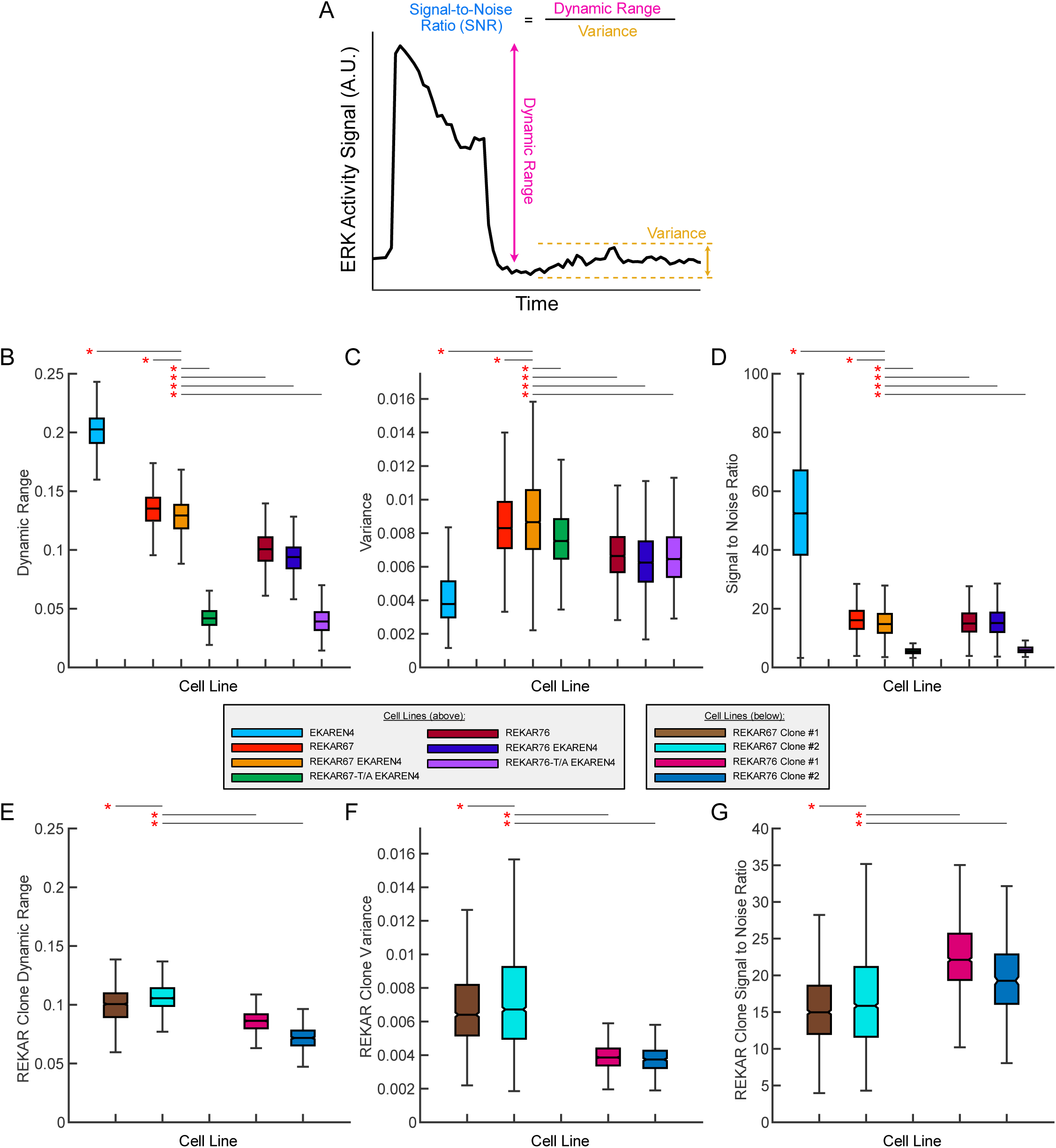
Activity trace characteristics and derivations. **(A)** Example plot indicating dynamic range (pink) as the distance between maximum and minimum signal, variance (yellow) as calculated using the post-inhibition signal, and Signal-to-Noise Ratio (SNR) (blue) as the quotient of the dynamic range and variance. **(B-D)** Box and Whisker plots showing the **(B)** dynamic range, **(C)** variance, and **(D)** SNR of EKAREN4, REKAR67, and REKAR76 expressing cells as indicated. **(E-G)** Box and whiskers plots showing REKAR clone’s **(E)** dynamic range, **(F)** signal variance, and **(G)** SNR, as indicated. Red asterisks indicate P < 0.0005 between the compared groups as determined by 1-way ANOVA adjusted for false-discovery rate using unnett’s multi-comparison adjustment.

In cell populations, we found that REKAR67 had a significantly larger Signal than REKAR76 (0.134 and 0.100 A.U.; P < 0.0005). Comparatively, EKAREN4 had a larger Signal value of 0.199. As expected, T/A mutants had very low Signal values of <0.05 (**Figure 3B**). REKAR67 also had significantly larger Noise than REKAR76 (0.00878 and 0.00691 A.U.; P < 0.0005) (**Figure 3C**). From these Signal and Noise values, we then calculated the SNR ratio for each reporter. In these calculations, the higher values of both Signal and Noise for REKAR67 offset each other, such that the two REKAR configurations were comparable in SNR (16.4 and 15.5; P < 0.0005). A similar calculation for EKAREN4 displayed a ∼3 fold larger SNR than either REKAR variant (EKAREN4 SNR = 53.3), due to its greater Signal and lower Noise measurements (**Figure 3D**).

Analysis of clonal populations of REKAR-expressing cells resulted in similar conclusions. Both clones of REKAR67 had higher Signal values (0.099122, 0.1065) than those of REKAR76 (0.0857, 0.0722) (**Figure 3E**). Similarly, REKAR67 clones had higher Noise (0.00702, 0.00754) than REKAR76 clones (0.00399, 0.00386) (**Figure 3F**). SNR values for clones of REKAR67 were 15.8 and 16.8, in comparison to 22.5 and 19.7 for REKAR76 (**Figure 3G**). Thus, REKAR76 appears to display a consistently greater SNR in these clonal populations.

## Discussion

The two new red/far red-FRET reported here add new capabilities to the existing toolbox of signaling biosensors. We demonstrate that these new constructs are essentially equivalent to existing ERK FRET biosensors in their cellular responses and can be easily multiplexed with green- or yellow-range biosensors (e.g., CFP/YFP or GFP-based biosensors). This capability will make it possible to measure ERK activity in tandem with the many cpGFP-based sensors now available for metabolites (including ATP, glucose, lactate, pyruvate, 1,6 bisphosphate [25–29], signaling intermediates and cofactors [30–32], and other kinases [13,17,33,34]. Another red-shifted variant of ERK reporter is also available, termed RAB-EKARev [34,35]. This biosensor is based on a similar design, but relies on dimerization-dependent fluorescence of ddRFP, rather than FRET. Overall, this reporter appears to have similar kinetic responses to growth factor stimulation, but its reversibility following ERK inhibition has not been established, and it was not directly compared to existing ERK biosensors. Additionally, the excitation and emission profile for REKAR67/76 is substantially further red-shifted relative to ddRFP. It should be possible in principle to multiplex CFP/YFP, ddRFP, and miRFP670/720 FRET-based reporters without significant spectral overlap.

While it was previously possible to obtain a readout of ERK activity in the red or far red range using ERK kinase translocation reporters (ERK-KTRs) [36], in which any color FP can be used, this approach requires compromises in the specificity of the reporter and the ability to monitor subcellular activity localization. Like the initial versions of EKAR, ERK-KTRs have some responsiveness to CDK activity [37], although a modified version may reduce this tendency [38]. The CDK responsiveness of ERK-KTR can be corrected mathematically when CDK2 biosensor measurements are also available [39], but this configuration adds substantial complexity to the imaging and data analysis. Moreover, KTRs are not able to provide measurements of kinase activity within specific subcellular regions, as they rely on changes in subcellular localization as their readout mode. Finally, KTR-based reporters can be problematic in situations where cell shapes are irregular or are changing during the experiment, as these changes can affect the ability to reliably sample and segment the cytoplasm without artifacts. Avoiding such artifacts will likely be particularly useful when imaging cells within tissue contexts, whether *in vivo* or *ex vivo*. We expect the REKAR biosensors to be particularly useful in these contexts, especially given the greater penetrance of red wavelengths through tissue.

With the limitations overcome by REKAR67 or REKAR76, it should be possible to design many new experiments with paired measurements for multiple kinases. Interestingly, REKAR67 or REKAR76 also make it possible to monitor ERK activity at different cellular locations simultaneously, because they can be paired with a differentially localized CFP/YFP ERK reporter. For example, ERK activity at the plasma membrane and in the nucleus, which have been reported to be distinct [24,40,41], could now be directly correlated within the same cell. While both EKAR and REKAR variants were localized to the nucleus in the dual-reporter experiments shown here, it would be straightforward to add alternate localization motifs to one or both reporters.

This work suggests a number of new questions to be addressed. First, the fact that both we and others obtained functional reporters through a simple swap in FPs without the need for substantial optimization, suggests that this strategy can be applied more broadly to existing FRET reporters. Additional work will be needed to determine what limits, if any, apply to such designs. Given that the structure of the miRFPs differs substantially from the beta barrel CFP and YFP structure, there may be important additional factors, such as linker length, to be further optimized. The smaller size of the miRFP fluorophores could change the way that the biosensor interacts with both kinases and phosphatases, and a more careful examination of how these effects on- and off-rates is warranted. Furthermore, the optical properties of miRFP-based FRET reporters will need to be experimentally determined, including the pH sensitivity of the readout, and the fluorescence lifetime characteristics.

Finally, we note some limitations in the REKAR sensors. Most notably, they appear to be subject to a greater degree of measurement noise. We have not fully pinpointed the source of this noise, but we suspect that it may arise in part from the imaging configuration we used, in which the same excitation wavelength is used to generate both the donor and acceptor images. Relative to the configuration used for the CFP/YFP biosensor, this setup results in an activity calculation that is more subject to spectral cross talk, which may amplify noise. Furthermore, imaging the emission wavelengths for both miRFP fluorophores simultaneously using a dual-camera configuration could reduce noise. Additionally, the miRFP fluorophores require biliverdin as a cofactor. While we found that it is optimal to add exogenous biliverdin, as in the experiments shown here, we also observed sufficient fluorescence without this addition, likely due to trace amounts of biliverdin available in the serum-containing growth medium during cell plating. Because biliverdin availability could change depending on the serum source or washing procedures used, it is an important variable to control to ensure experimental replicability.

**Supplemental Figure 1.**
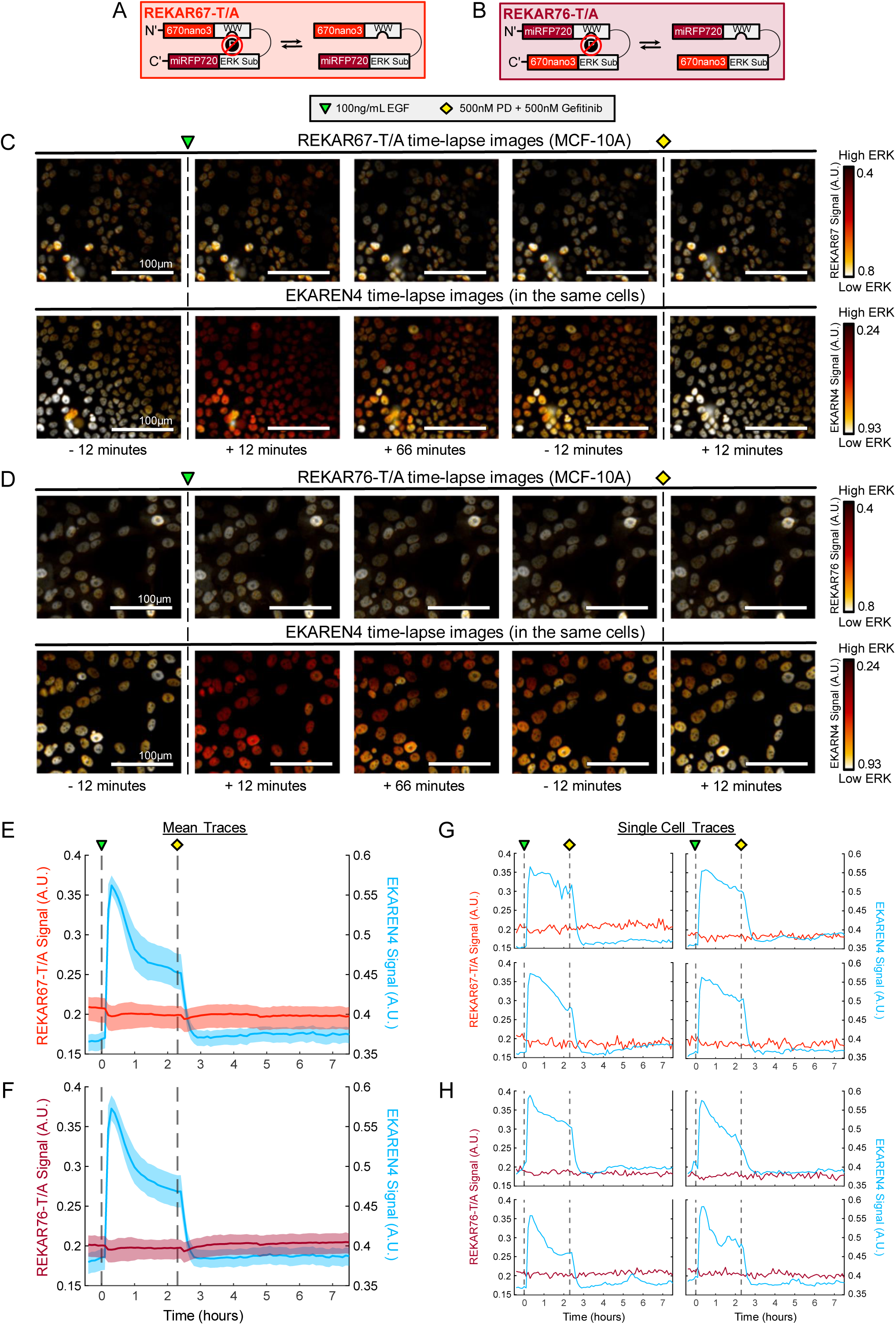
Non-functional REKAR Control Designs and Validation. **(A-B)** Graphical representations of the mutated ERK biosensors: REKAR67 T/A, and REKAR76 T/A. **(C-D)** *Top row* shows REKAR67 or REKAR76 and *bottom row* shows EKAREN4 pseudo-colored images indicating the relative ERK activity in MCF10A cells coexpressing the REKAR and EKAREN4 biosensors (white = low ERK, red = high ERK). Images show before and after EGF stimulation then MEK/EGFR inhibition. Scale bars are 100μm. **(E-F)** Mean FRET signal of REKAR67 (red) or REKAR76 (maroon) overlaid on the mean EKAREN4 signal (blue) of the same cells. **(G-H)** Normalized data from **E&F**.

## SUPPLEMENTAL VIDEO LEGENDS

**Supplementary Video 1:** MCF-10A cells that simultaneously express both REKAR67 and EKAREN4 ERK biosensors. The movie shows pseudo-colored images made by taking the ratio of miRFP670nano3 and FRET720 channels (REKAR67). Cells are treated with 100 ng/mL EGF 30 minutes into the movie. Cells are also treated with 500 nM PD-0325901 and 500 nM Gefitinib 2.5 hours into the movie. Single frames of this movie were used to make Figure 1 of the paper.

**Supplementary Video 2:** MCF-10A cells that simultaneously express both REKAR67 and EKAREN4 ERK biosensors. The movie shows pseudo-colored images made by taking the ratio of CFP and YFP channels (EKAREN4). Cells are treated with 100 ng/mL EGF 30 minutes into the movie. Cells are also treated with 500 nM PD-0325901 and 500 nM Gefitinib 2.5 hours into the movie. Single frames of this movie were used to make Figure 1 of the paper.

**Supplementary Video 3:** MCF-10A cells that simultaneously express both REKAR76 and EKAREN4 ERK biosensors. The movie shows pseudo-colored images made by taking the ratio of miRFP670nano3 and FRET720 channels (REKAR76). Cells are treated with 100 ng/mL EGF 30 minutes into the movie. Cells are also treated with 500 nM PD-0325901 and 500 nM Gefitinib 2.5 hours into the movie. Single frames of this movie were used to make Figure 1 of the paper.

**Supplementary Video 4:** MCF-10A cells that simultaneously express both REKAR76 and EKAREN4 ERK biosensors. The movie shows pseudo-colored images made by taking the ratio of CFP and YFP channels (EKAREN4). Cells are treated with 100 ng/mL EGF 30 minutes into the movie. Cells are also treated with 500 nM PD-0325901 and 500 nM Gefitinib 2.5 hours into the movie. Single frames of this movie were used to make Figure 1 of the paper.

**Supplementary Video 5:** MCF-10A cells that simultaneously express both REKAR67-T/A and EKAREN4 ERK biosensors. The movie shows pseudo-colored images made by taking the ratio of miRFP670nano3 and FRET720 channels (REKAR67-T/A). Cells are treated with 100 ng/mL EGF 30 minutes into the movie. Cells are also treated with 500 nM PD-0325901 and 500 nM Gefitinib 2.5 hours into the movie. Single frames of this movie were used to make Supplementary Figure 1 of the paper.

**Supplementary Video 6:** MCF-10A cells that simultaneously express both REKAR67-T/A and EKAREN4 ERK biosensors. The movie shows pseudo-colored images made by taking the ratio of CFP and YFP channels (EKAREN4). Cells are treated with 100 ng/mL EGF 30 minutes into the movie. Cells are also treated with 500 nM PD-0325901 and 500 nM Gefitinib 2.5 hours into the movie. Single frames of this movie were used to make Supplementary Figure 1 of the paper.

**Supplementary Video 7:** MCF-10A cells that simultaneously express both REKAR76-T/A and EKAREN4 ERK biosensors. The movie shows pseudo-colored images made by taking the ratio of miRFP670nano3 and FRET720 channels (REKAR76-T/A). Cells are treated with 100 ng/mL EGF 30 minutes into the movie. Cells are also treated with 500 nM PD-0325901 and 500 nM Gefitinib 2.5 hours into the movie. Single frames of this movie were used to make Supplementary Figure 1 of the paper.

**Supplementary Video 8:** MCF-10A cells that simultaneously express both REKAR76-T/A and EKAREN4 ERK biosensors. The movie shows pseudo-colored images made by taking the ratio of CFP and YFP channels (EKAREN4). Cells are treated with 100 ng/mL EGF 30 minutes into the movie. Cells are also treated with 500 nM PD-0325901 and 500 nM Gefitinib 2.5 hours into the movie. Single frames of this movie were used to make Supplementary Figure 1 of the paper.

## Author Contributions

Conceptualization and design: NLD, JGA

Study data collection: NLD, JYH, FX, BR

Raw image data analysis: NLD, JYH, FX

Programming/Coding: NLD, JYH, BR, MP

Results Interpretation: All authors

Drafting/Writing/Editing the manuscript: All authors

Figure creation / revision: NLD, JYH, FX

## Funding Sources

<u>National Heart, Lung, and Blood Institute (NHLBI):</u>

HL007013 (NLD) (UC Davis Lung Center T32)

R01 HL151983-01A1 (JGA)

<u>National Institute of General Medical Sciences (NIGMS):</u>

R35GM139621 (JGA)

## References

1. Ram, A.; Murphy, D.; DeCuzzi, N.; Patankar, M.; Hu, J.; Pargett, M.; Albeck, J.G. A Guide to ERK Dynamics, Part 1: Mechanisms and Models. Biochem. J 2023, 480, 1887–1907.

2. Ram, A.; Murphy, D.; DeCuzzi, N.; Patankar, M.; Hu, J.; Pargett, M.; Albeck, J.G. A Guide to ERK Dynamics, Part 2: Downstream Decoding. Biochem. J 2023, 480, 1909–1928.

3. Nakamura, A.; Goto, Y.; Kondo, Y.; Aoki, K. Shedding Light on Developmental ERK Signaling with Genetically Encoded Biosensors. Development 2021, 148, doi:10.1242/dev.199767.

4. Hiratsuka, T.; Fujita, Y.; Naoki, H.; Aoki, K.; Kamioka, Y.; Matsuda, M. Intercellular Propagation of Extracellular Signal-Regulated Kinase Activation Revealed by in Vivo Imaging of Mouse Skin. Elife 2015, 4, e05178.

5. DeCuzzi, N.L.; Oberbauer, D.; Chmiel, K.J.; Pargett, M.; Ferguson, J.M.; Murphy, D.; Hardy, M.; Ram, A.; Zeki, A.A.; Albeck, J.G. Spatiotemporal Clusters of ERK Activity Coordinate Cytokine-Induced Inflammatory Responses in Human Airway Epithelial Cells. Am. J. Respir. Cell Mol. Biol. 2024, doi:10.1165/rcmb.2024-0256OC.

6. Ender, P.; Gagliardi, P.A.; Gagliardi, M.; Dobrzyński, A.; Fristmantiene, A.; Dessauges, C.; Hӧhener, T.; Spatiotemporal Control of ERK Pulse Frequency Coordinates Fate Decisions during Mammary Acinar Morphogenesis. Dev. Cell 2022, 57, 2153–2167.e6.

7. Gagliardi, .A.; Dobryński, M.; Jacques, M.-A.; Dessauges, C.; Ender, P.; Blum, Y.; Hughes, R.M.; Cohen, A.R.; Pertz, O. Collective ERK/Akt Activity Waves Orchestrate Epithelial Homeostasis by Driving Apoptosis-Induced Survival. Dev. Cell 2021, doi:10.1016/j.devcel.2021.05.007.

8. Valon, L.; Davidović, A.; Levillayer, F.; Villars, A.; Chouly, M.; Cerqueira-Campos, F.; Levayer, R. Robustness of Epithelial Sealing Is an Emerging Property of Local ERK Feedback Driven by Cell Elimination. Dev. Cell 2021, 56, 1700–1711.e8.

9. Aikin, T.J.; Peterson, A.F.; Pokrass, M.J.; Clark, H.R.; Regot, S. MAPK Activity Dynamics Regulate Non-Cell Autonomous Effects of Oncogene Expression. Elife 2020, 9, doi:10.7554/eLife.60541.

10. Gillies, T.E.; Pargett, M.; Silva, J.M.; Teragawa, C.K.; McCormick, F.; Albeck, J.G. Oncogenic Mutant RAS Signaling Activity Is Rescaled by the ERK/MAPK Pathway. Mol. Syst. Biol. 2020, 16, e9518.

11. Davies, A.E.; Pargett, M.; Siebert, S.; Gillies, T.E.; Choi, Y.; Tobin, S.J.; Ram, A.R.; Murthy, V.; Juliano, C.; Quon, G.;, et al. Systems-Level Properties of EGFR-RAS-ERK Signaling Amplify Local Signals to Generate Dynamic Gene Expression Heterogeneity. Cell Syst 2020, 11, 161–175.e5.

12. Harvey, C.D.; Ehrhardt, A.G.; Cellurale, C.; Zhong, H.; Yasuda, R.; Davis, R.J.; Svoboda, K. A Genetically Encoded Fluorescent Sensor of ERK Activity. Proc. Natl. Acad. Sci. U. S. A. 2008, 105, 19264–19269.

13. Komatsu, N.; Aoki, K.; Yamada, M.; Yukinaga, H.; Fujita, Y.; Kamioka, Y.; Matsuda, M. Development of an Optimized Backbone of FRET Biosensors for Kinases and GTPases. Mol. Biol. Cell 2011, 22, 4647–4656.

14. Vandame, P.; Spriet, C.; Riquet, F.; Trinel, D.; Cailliau-Maggio, K.; Bodart, J.-F. Optimization of ERK Activity Biosensors for Both Ratiometric and Lifetime FRET Measurements. Sensors 2014, 14, 1140–1154.

15. Ponsioen, B.; Post, J.B.; Buissant des Amorie, J.R.; Laskaris, D.; van Ineveld, R.L.; Kersten, S.; Bertotti, A.; Sassi, F.; Sipieter, F.; Cappe, B.;, et al. Quantifying Single-Cell ERK Dynamics in Colorectal Cancer Organoids Reveals EGFR as an Amplifier of Oncogenic MAPK Pathway Signalling. Nat. Cell Biol. 2021, 23, 377–390.

16. Chavez-Abiega, S.; Grönloh, M.L.B.; Gadella, T.W.J.; Bruggeman, F.J.; Goedhart, J. Single-Cell Imaging of ERK and Akt Activation Dynamics and Heterogeneity Induced by G-Protein-Coupled Receptors. J. Cell Sci. 2022, 135, doi:10.1242/jcs.259685.

17. Chen, M.; Sun, T.; Zhong, Y.; Zhou, X.; Zhang, J. A Highly Sensitive Fluorescent Akt Biosensor Reveals Lysosome-Selective Regulation of Lipid Second Messengers and Kinase Activity. ACS Cent. Sci. 2021, doi:10.1021/acscentsci.1c00919.

18. Pargett, M.; Gillies, T.E.; Teragawa, C.K.; Sparta, B.; Albeck, J.G. Single-Cell Imaging of ERK Signaling Using Fluorescent Biosensors. In Kinase Signaling Networks; Tan, A.-C., Huang, P.H., Eds.; Springer New York: New York, NY, 2017; pp. 35–59 ISBN 9781493971541.

19. Pargett, M.; Albeck, J.G. Live-Cell Imaging and Analysis with Multiple Genetically Encoded Reporters. Curr. Protoc. Cell Biol. 2018, 78, 4.36.1–4.36.19.

20. Debnath, J.; Muthuswamy, S.K.; Brugge, J.S. Morphogenesis and Oncogenesis of MCF-10A Mammary Epithelial Acini Grown in Three-Dimensional Basement Membrane Cultures. Methods 2003, 30, 256–268.

21. Sparta, B.; Pargett, M.; Minguet, M.; Distor, K.; Bell, G.; Albeck, J.G. Receptor Level Mechanisms Are Required for Epidermal Growth Factor (EGF)-Stimulated Extracellular Signal-Regulated Kinase (ERK) Activity Pulses. J. Biol. Chem. 2015, 290, 24784–24792.

22. Oliinyk, O.S.; Baloban, M.; Clark, C.L.; Carey, E.; Pletnev, S.; Nimmerjahn, A.; Verkhusha, V.V. Single-Domain near-Infrared Protein Provides a Scaffold for Antigen-Dependent Fluorescent Nanobodies. Nat. Methods 2022, 19, 740–750.

23. Shcherbakova, D.M.; Cox Cammer, N.; Huisman, T.M.; Verkhusha, V.V.; Hodgson, L. Direct Multiplex Imaging and Optogenetics of Rho GTPases Enabled by near-Infrared FRET. Nat. Chem. Biol. 2018, 14, 591–600.

24. Keyes, J.; Ganesan, A.; Molinar-Inglis, O.; Hamidzadeh, A.; Zhang, J.; Ling, M.; Trejo, J.; Levchenko, A.; Zhang, J. Signaling Diversity Enabled by Rap1-Regulated Plasma Membrane ERK with Distinct Temporal Dynamics. Elife 2020, 9, doi:10.7554/eLife.57410.

25. Tantama, M.; Martínez-François, J.R.; Mongeon, R.; Yellen, G. Imaging Energy Status in Live Cells with a Fluorescent Biosensor of the Intracellular ATP-to-ADP Ratio. Nat. Commun. 2013, 4, 2550.

26. San Martín, A.; Ceballo, S.; Ruminot, I.; Lerchundi, R.; Frommer, W.B.; Barros, L.F. A Genetically Encoded FRET Lactate Sensor and Its Use to Detect the Warburg Effect in Single Cancer Cells. PLoS One 2013, 8, e57712.

27. Kondo, H.; Ratcliffe, C.D.H.; Hooper, S.; Ellis, J.; MacRae, J.I.; Hennequart, M.; Dunsby, C.W.; Anderson, K.I.; Sahai, E. Single-Cell Resolved Imaging Reveals Intra-Tumor Heterogeneity in Glycolysis, Transitions between Metabolic States, and Their Regulatory Mechanisms. Cell Rep. 2021, 34, 108750.

28. Koberstein, J.N.; Stewart, M.L.; Smith, C.B.; Tarasov, A.I.; Ashcroft, F.M.; Stork, P.J.S.; Goodman, R.H. Monitoring Glycolytic Dynamics in Single Cells Using a Fluorescent Biosensor for Fructose 1,6-Bisphosphate. Proc. Natl. Acad. Sci. U. S. A. 2022, 119, e2204407119.

29. Arce-Molina, R.; Cortés-Molina, F.; Sandoval, P.Y.; Galaz, A.; Alegría, K.; Schirmeier, S.; Barros, L.F.; San Martín, A. A Highly Responsive Pyruvate Sensor Reveals Pathway-Regulatory Role of the Mitochondrial Pyruvate Carrier MPC. Elife 2020, 9, e53917.

30. Park, J.G.; Palmer, A.E. Quantitative Measurement of Ca2+ and Zn2+ in Mammalian Cells Using Genetically Encoded Fluorescent Biosensors. Methods Mol. Biol. 2014, 1071, 29–47.

31. Tian, L.; Hires, S.A.; Mao, T.; Huber, D.; Chiappe, M.E.; Chalasani, S.H.; Petreanu, L.; Akerboom, J.; McKinney, S.A.; Schreiter, E.R.;, et al. Imaging Neural Activity in Worms, Flies and Mice with Improved GCaMP Calcium Indicators. Nat. Methods 2009, 6, 875–881.

32. Wang, L.; Wu, C.; Peng, W.; Zhou, Z.; Zeng, J.; Li, X.; Yang, Y.; Yu, S.; Zou, Y.; Huang, M.;, et al. A High-Performance Genetically Encoded Fluorescent Indicator for in Vivo cAMP Imaging. Nat. Commun. 2022, 13, 5363.

33. Schmitt, D.L.; Curtis, S.D.; Lyons, A.C.; Zhang, J.-F.; Chen, M.; He, C.Y.; Mehta, S.; Shaw, R.J.; Zhang, J. Spatial Regulation of AMPK Signaling Revealed by a Sensitive Kinase Activity Reporter. Nat. Commun. 2022, 13, 3856.

34. Mehta, S.; Zhang, Y.; Roth, R.H.; Zhang, J.-F.; Mo, A.; Tenner, B.; Huganir, R.L.; Zhang, J. Single-Fluorophore Biosensors for Sensitive and Multiplexed Detection of Signalling Activities. Nat. Cell Biol. 2018, 20, 1215–1225.

35. Ding, Y.; Li, J.; Enterina, J.R.; Shen, Y.; Zhang, I.; Tewson, P.H.; Mo, G.C.H.; Zhang, J.; Quinn, A.M.; Hughes, T.E.;, et al. Ratiometric Biosensors Based on Dimerization-Dependent Fluorescent Protein Exchange. Nat. Methods 2015, 12, 195–198.

36. Regot, S.; Hughey, J.J.; Bajar, B.T.; Carrasco, S.; Covert, M.W. High-Sensitivity Measurements of Multiple Kinase Activities in Live Single Cells. Cell 2014, 157, 1724– 1734.

37. Gerosa, L.; Chidley, C.; Fröhlich, F.; Sanchez, G.; Lim, S.K.; Muhlich, J.; Chen, J.-Y.; Vallabhaneni, S.; Baker, G.J.; Schapiro, D.;, et al. Receptor-Driven ERK Pulses Reconfigure MAPK Signaling and Enable Persistence of Drug-Adapted BRAF-Mutant Melanoma Cells. Cell Syst 2020, 11, 478–494.e9.

38. Wilcockson, S.G.; Guglielmi, L.; Araguas Rodriguez, P.; Amoyel, M.; Hill, C.S. An Improved Erk Biosensor Detects Oscillatory Erk Dynamics Driven by Mitotic Erasure during Early Development. Dev. Cell 2023, 58, 2802–2818.e5.

39. Kim, S.; Carvajal, R.; Kim, M.; Yang, H.W. Kinetics of RTK Activation Determine ERK Reactivation and Resistance to Dual BRAF/MEK Inhibition in Melanoma. Cell Rep. 2023, 42, 112570.

40. Cohen-Saidon, C.; Cohen, A.A.; Sigal, A.; Liron, Y.; Alon, U. Dynamics and Variability of ERK2 Response to EGF in Individual Living Cells. Mol. Cell 2009, 36, 885–893.

41. Nakakuki, T.; Birtwistle, M.R.; Saeki, Y.; Yumoto, N.; Ide, K.; Nagashima, T.; Brusch, L.; Ogunnaike, B.A.; Okada-Hatakeyama, M.; Kholodenko, B.N. Ligand-Specific c-Fos Expression Emerges from the Spatiotemporal Control of ErbB Network Dynamics. Cell 2010, 141, 884–896.

